# Complex genotype-phenotype relationships shape the response to treatment of Down Syndrome Childhood Acute Lymphoblastic Leukaemia

**DOI:** 10.1101/2022.02.06.479302

**Authors:** Christoph Lutz, Virginia A. Turati, Ruth Clifford, Petter S. Woll, Thomas Stiehl, Anders Castor, Sally A. Clark, Helen Ferry, Veronica Buckle, Andreas Trumpp, Anthony Ho, Anna Marciniak-Czochra, Javier Herrero, Anna Schuh, Sten-Eirik W. Jacobsen, Tariq Enver

## Abstract

Extensive genetic and epigenetic variegation has been demonstrated in many malignancies. Importantly, their interplay has the potential to contribute to disease progression and treatment resistance. To shed light on the complex relationships between these different sources of intra-tumour heterogeneity, we explored their relative contributions to the evolutionary dynamics of Acute Lymphoblastic Leukemia (ALL) in children with Down syndrome, which has particularly poor prognosis. We quantified the tumour propagating potential of genetically distinct sub-clones using serial transplantation assays and SNP-arrays. While most leukaemias were characterized by a single dominant subclone, others were highly heterogeneous. Importantly, we provide clear and direct evidence that genotypes and phenotypes with functional relevance to leukemic progression and treatment resistance can co-segregate within the disease. Hence, individual genetic lesions can be restricted to well-defined cell immunophenotypes, corresponding to different stages of the leukemic differentiation hierarchy and varied proliferation potentials. As a result of this difference in fitness, which can be accurately quantified via competitive transplantation assays, matching diagnostic, post-treatment, and relapse leukemias can be dominated by different genotypes, including pre-leukemic clones persisting throughout the disease progression and treatment. Intriguingly, plasticity also appears to be a temporally defined property that can segregate with genotype. These results suggest that Down Syndrome ALL should be viewed as a complex matrix of cells exhibiting genetic and epigenetic heterogeneity that foster extensive clonal evolution and competition. Therapeutic intervention reshapes this ‘eco-system’ and may provide the right conditions for the preferential expansion of selected compartments and subsequently relapse.

## Introduction

The existence of intra-tumour genetic heterogeneity, both genetic and epigenetic, has been documented in both solid and haematological malignancies^1–3^. In studies of childhood acute lymphoblastic leukaemia, we and others previously used multicolour-FISH and whole genome sequencing to show that these tumours are clonally variegated, and to demonstrate through serial transplantation assays, that leukaemia tumour propagating cells (TPC) are also genetically heterogeneous^4, 5^. This finding was supported by our later genetic analysis of immunophenotypically distinct ALL diagnostic subpopulations which demonstrated similar distribution of subclones within a disease-specific primitive B-cell compartment (StemB, 34^+^38^-^19^+^) and the disease bulk in most, but, crucially, not all leukaemias^6^. Such experiments relied on high-depth bulk whole-genome sequencing followed by single-cell qPCR and, together with our earlier work, suggested that genotype and phenotypes related to differentiation stage and cell-cycle state might not necessarily co-segregate within the disease. Importantly, as chemotherapy-induced selection acts on these same phenotypic traits^6–8^, this lack of segregation also underpinned the survival of most genotypes to the early stages of childhood ALL treatment (namely, induction chemotherapy – where maximum cytotoxicity is achieved); as we demonstrated using a xenograft model of transplantation and treatment of primary leukaemias^6^.

Large-scale comparisons of matching diagnostic and relapse samples from paediatric ALL patients, have nonetheless previously reported that in some patients a subset of diagnostic subclones, or of their later evolutionary descendants, can be selectively enriched at the time of disease recurrence^9, 10^. Assuming that these clones did not become dominant as a result of stochastic selection, this finding opens to the possibility that fitness-based genotypic selection might potentially occur on a longer timescale than that of induction chemotherapy. It is not unreasonable to speculate that in a treatment protocol that lasts up to four years even subtle differences in fitness might eventually lead to clonal dominance. Another possibility is that in certain settings, genotype, and treatment-associated phenotypes do, in fact, segregate (either already at diagnosis or later because of further genomic evolution). In support of this latter hypothesis, our analysis of 34^+^38^-^19^+^ (StemB) ALL cells indicated that in a small proportion of leukaemias (1 out of 5 analysed), specific genotypes might indeed be preferentially enriched within developmentally more primitive and quiescent disease compartments^6^. Furthermore, recent evidence suggests thar mutations induced by chemotherapy treatment itself (particularly those targeting p53) might be responsible for - post-diagnosis – generating new, fitter subclones, and inducing drug-resistance; therefore, leading to relapse^11^.

To shed light on the complex genotype-phenotype relationship that shape ALL, we here conduct a comprehensive investigation into the relative contribution of different sources of intratumor heterogeneity, namely genetic makeup and epigenetically determined properties - particularly cell cycle state, TPC activity and differentiation stage - to the disease initiation, sustenance, and treatment resistance. In doing so we focused our attention Down’s syndrome associated ALL as this rarer disease subtype has so far remained poorly characterised while being associated with considerably worse prognosis than ALL in children without Down syndrome^12, 13^.

Altogether our results indicate that ALL should be viewed as a complex matrix of cells exhibiting genetic and epigenetic heterogeneity. Our work provides evidence for, in some patients, diversification in functional properties amongst immunophenotypically-distinct leukemic subpopulations. In these setting, those rare leukemic cells which remain after induction chemotherapy, and primarily account for minimal residual disease (MRD), are highly enriched for the stem/B phenotype; suggesting that the more primitive and less proliferative leukemic compartments likely constitute the pool of cells from which relapse arises. Crucially, in this setting where genotype and resistance-associated phenotypes co-segregate, bottleneck selection at phenotype level is accompanied by evidence of a clonal sweep.

## Methods

### Patient samples

Diagnostic and follow-up ALL bone marrow (BM) samples were obtained after informed consent and the approval of the relevant research ethics committee from patients at the Paediatrics Haematology Unit, Lund University Hospital, Sweden. Samples from 4 children with Down’s ALL were analyzed. For further patient characteristics see Table 1.

### Cell separation, phenotyping, and sorting

Total mononuclear cells (MNCs) were isolated by ficoll gradient centrifugation and directly cryopreserved in DMSO for later use. In some cases, CD34^+^ cells were enriched by magnetic bead separation (StemCell Technologies or Miltenyi). After thawing, dead cells were evaluated and excluded by FACS after staining with DAPI. CD34-enriched cells or MNCs from BM were stained with anti-CD19 PE (BD-Pharmingen), CD34 FITC (BD-Pharmingen) and CD38 APC (BD-Pharmingen). Cells were analyzed in staining buffer containing DAPI at 0·1µg/ml, for live cell analysis. I.) HSC (34^+^38^-^19^-^); II.) Stem/B (34^+^38^-^ 19^+^) III.) Progenitor (34^+^38^+^19^-^) IV.) ProB (34^+^38^+^19^+^) and V.) PreB/Mature B-cells (34^-^19^+^) were purified by flow cytometry (FACS Aria, BD-Pharmingen). Data acquisition and analysis were done with CellQuest (Beckon Dickinson) or FlowJo (Tree star) software.

### Cell cycle analysis of leukaemia subpopulations

Cells were stained with CD19-PECy5 (BioLegend), then fixed and permeabilized with 1.6% paraformaldehyde and 90% ice-cold methanol. This was followed by staining with CD34-APC (BD), CD38-PETxR (Invitrogen) and Ki67-FITC (Becton-Dickinson). For DNA content analysis cells were incubated with 0·5µg/mL DAPI. Stained cells were analyzed by excitation of DAPI with a violet laser on a FACS LSRII SORP (Becton-Dickinson) at the University of Oxford (UK).

### FISH analysis

Cells were fixed on slides in methanol/acetic acid fixative (3:1 vol/vol) and then hybridized with BCR-ABL1 FISH probes using the LSI BCR/ABL1 Dual Color Dual Fusion Translocation probes (Abbot). The slides were analyzed with an Olympus BX51 microscope equipped with epifluorescence and a triple band pass filter. Images were captured by using a Sensys charge-coupled device camera (Photometrics, Tucson, AZ) and MacProbe software (Applied Imaging, Newcastle upon Tyne, U.K.). When possible, and unless otherwise specified at least 100-150 nuclei were analyzed in each sample

### SNP-Array

SNP-Arrays we performed as previously describe in Schuh et al., 2012^14^

### NSG-mouse transplantation assay

Primary childhood ALL cells were transplanted into 8-12 weeks old NOD/SCID IL2Rγ^hull^ (NSG) sub-lethally irradiated mice (either females or males) via intramedullary injection. To minimize possible adverse effects of sublethal irradiation, mice were administered acid water for a week prior to the procedure, and Baytril (resuspended at 25.5 mg/kg in the drinking water) for the 2 weeks following it. Sub-lethal irradiation was achieved with a single dose of 375 cGy. Unless otherwise stated, each mouse received 2 x 10^5^ primary leukemia cells resuspended in 40μl PBS 0.5% FBS. In the case of secondary limiting dilution assays, a specified equal dose of treated and control leukemic cells harvested from the BM of primary recipients was injected. Twelve-week post-injection mice were sampled by bone marrow aspiration and the percentage of human engraftment was evaluated by flow cytometry (hCD45/(hCD45^+^ + mCD45^+^)). At the same time, human cells were also FACS sorted for downstream applications. Mice displaying at least 70% human engraftment were then randomly assigned to either control or treatment groups (see below for details on the treatment protocol). At the end of the treatment window all mice were culled, and their tibias, femurs, pelvises, spleen, and brain harvested, and cells were then stained for FACS sorting.

### Modeling of clonal evolution

See supplementary materials.

### Small cell numbers RNA sequencing

Equivalent cell numbers (400 cells per sample) were flow sorted directly into 800μl Trizol reagent (Invitrogen) and snap frozen in dry ice (long term storage at -80C). At the time of extraction, the samples were thawed at RT and 160ul of chloroform was added to each. Following a centrifugation step the RNA was isolated from the aqueous phase and precipitated through the addition of equal volumes of isopropanol supplemented with 20μg linear polyacrylamide. Samples were washed twice in 80% ethanol (first wash over night at 4°C, second wash 5 minutes at RT). RNA pellets were resuspended in 3-15μl of diethylpyrocarbonate treated water (DEPC). RNA was then quantified by loading of 0.5-1ul on an Agilent Bionalyser RNA 6,000 pico chip. Where possible equivalent amounts of total RNA (100pg) from all samples were used for first strand synthesis with the SmartERv3 kit (Takara Clontech) followed by 15-18 cycles of amplification (according to manufacturers’ instruction). cDNA was purified on Agencourt AMPureXP magnetic beads, washed twice with fresh 80% ethanol and eluted in 17μl elution buffer. 1μl cDNA was quantified with Qubit dsDNA HS (Molecular Probes) and checked on an Agilent Bioanalyser high sensitivity DNA chip. Sequencing libraries were produced from 150pg input cDNA using Illumina Nextera XT library preparation kit. A 1:4 miniaturized version of the protocol was adopted (see “Fluidigm Single-Cell cDNA Libraries for mRNA sequencing”, PN_100-7168_L1). Tagmentation time was 5mins, followed by 12 cycles of amplification using Illumina XT 24 or 96 index primer kit. Libraries were then pooled (1-2ul per sample depending on the total number of samples) and purified with equal volumes (1:1) of Agencourt AMPureXP magnetic beads. Final elution was in 66-144ul of resuspension buffer (depending on the total number of pooled samples). Libraries were checked on an Agilent Bioanalyser high sensitivity DNA chip (size range 150-2000bp) and quantified by Qubit dsDNA HS (Molecular Probes). Libraries were sequenced on Illumina® NextSeq 500 using 150bp paired end kits as per manufacturer’s instructions.

### Bioinformatics

Sequencing data was assessed to detect sequencing failures using FASTQC and lower quality reads were filtered or trimmed using *TrimGalore* (https://github.com/FelixKrueger/TrimGalore). Outlier samples containing low sequencing coverage or high duplication rates were discarded.

### Processing of bulk RNAseq

Bulk RNAseq samples were mapped to the human reference GRCh38 using *tophat2* (https://ccb.jhu.edu/software/tophat/index.shtml). Analyses were performed within the R statistical computing framework, version 3.5 using packages from *BioConductor version 3.7* (https://Bioconductor.org). Data was combined into a per-gene count matrix using featureCounts from the subread package. The *DEseq2* BioConductor package was used for outlier detection, normalization and differential gene expression analyses. All downstream analyses used Rlog transformed data.

### Data availability

SNP-array and RNAseq data are available at EGA under accession number EGAS00001005968.

## Results

We here explored the role of intra-tumour heterogeneity in the progression and response to treatment of Down’s associated ALL. We initially focused on genetic variegation and described the phylogenetic hierarchies of diagnostic material from 4 patients. Serial transplantation of bulk ALL samples into immune-deficient NSG-mice was concomitantly utilized to evaluate the tumour propagating cell activity (TPC) and dynamic behaviours of the different subclones identified. SNP-arrays were performed on diagnostic specimens as well as cells isolated from the BM of primary and secondary transplanted mice (see schemata in Figure 1A). In all patients, but one, a single dominant clone was detected both at diagnosis and in all primary and secondary transplanted mice (Figure 1B). In patient 2, instead, a more complex picture was observed, with several subclones coexisting within the tumour and contributing to a different extent to engraftment across recipients. In this patient we detected two deletions, on chromosomes 9 and 14 respectively, as well as an amplification on chromosome X (Figure 1C). Considering the mutational mosaicism, we observed in some of the engrafted mice, these observations allowed re-construction of a simple clonal phylogenetic architecture (Figure 1D, and Supp Material for Mosaicism data). However, a higher-depth genetic analysis of chromosome 9 suggested a far more complicated clonal landscape. A heterozygous chromosome 9 deletion was detected across engrafted animals with different degrees of homozygous involvement: to the extreme of copy neutral loss of heterozygosity (cnLOH) (Figure 2A). A simple explanation to this observation is that pre-existing sub-clones present at diagnosis at sub-threshold levels, were expanded through secondary transplantation until reaching detection levels. Alternatively, continuous broadening of the chromosome 9 deletion could have occurred over the short time frame of the serial transplantation assays. This latter hypothesis would be indicative of active ongoing clonal evolution. Based on the mosaicism data, and assuming no new mutations developed during transplantation, we were able to reconstruct a more comprehensive phylogenetic tree. To do so, we developed a custom-made mathematical model which we used to calculate all the theoretically possible diagnostic clone-combinations supported by our SNPs data analysis (Supplementary Table 1). In total 15-29 diagnostic clones were necessary to explain the mutational read-out from the xenografts. Of note, since SNP-arrays are limited in both genomic resolution (i.e., deletion size) and sensitivity to low abundance copy number variants (CNV) our model would if anything underestimate the number of clones (Figure 2B and supplementary methods). Nonetheless, this finding clearly illustrates the high clonal complexity of ALL and is line with our earlier single-cell WGS based characterization of genetic heterogeneity in non-Down Syndrome ALL, which also identified branching clonal architectures with up to 19 descendant subclones^6^.

**Figure 1:**
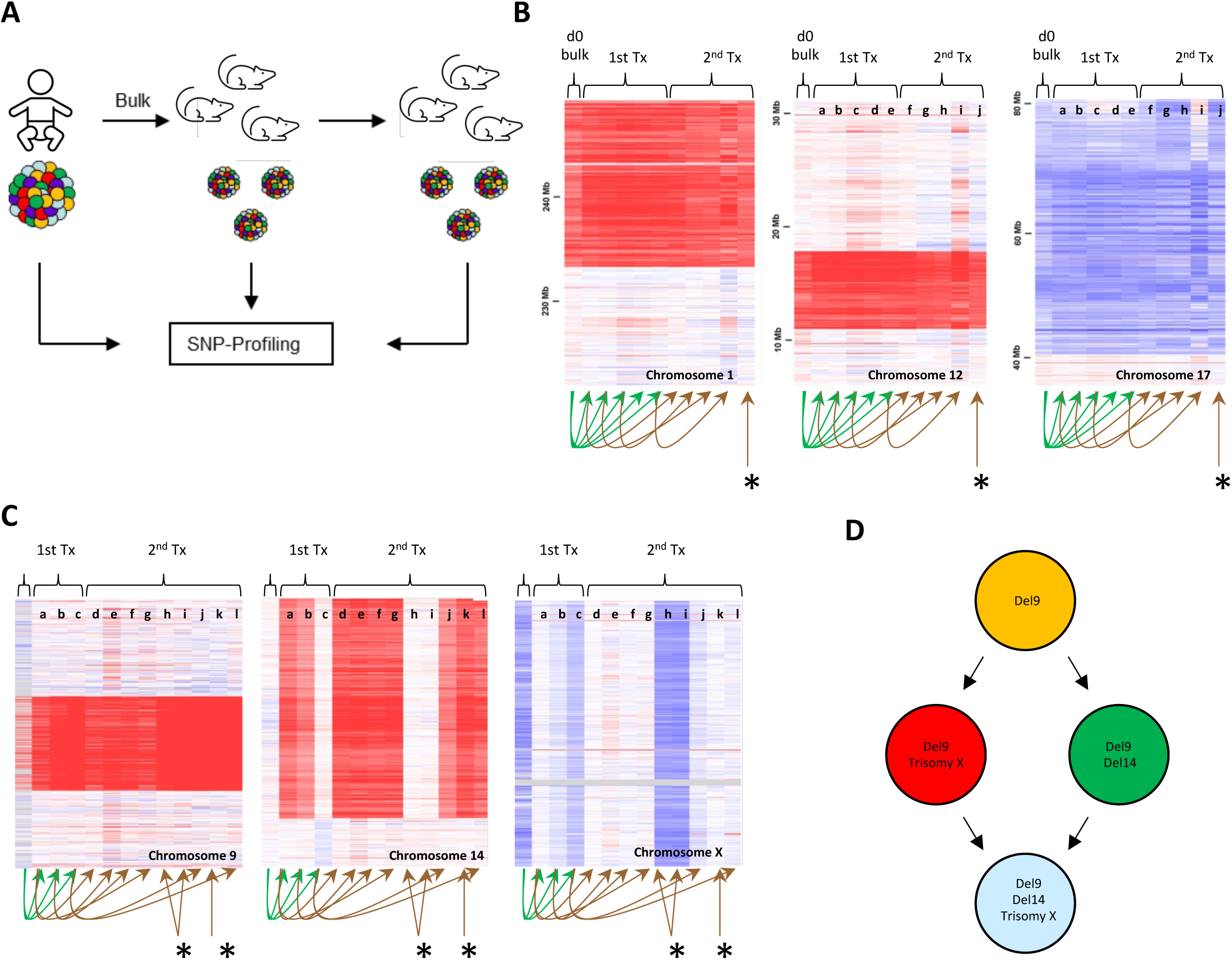
Study of clonal dynamics through serial transplantation assays. **A.** Experimental setup. BM samples before and after serial transplantation in NSG-mice were analyzed by SNP-arrays. **B.** SNP array results of patient 1. Deletions on Chromosomes 1, 12 and a single amplification on Chromosome 17 were identified and shown to be stable during serial transplantation. Relationships between transplants recipients are indicated with arrows. **C.** SNP array results of patient 2. Deletions on Chromosomes 9, 14 and a single amplification on Chromosome X were identified. During serial transplantation copy number abnormalities (CNA) disappeared and re-appeared suggesting the co-existence of different sub-clones. Relationships between transplants recipients are indicated with arrows. **D.** Initial reconstruction of the clonal hierarchy of patient 2 leukemia considering Chromosomes 9, 14 and X. * Indicate secondary transplants from a primary transplant for which SNP-array data are unavailable due to insufficient numbers of cells for analysis.

**Figure 2:**
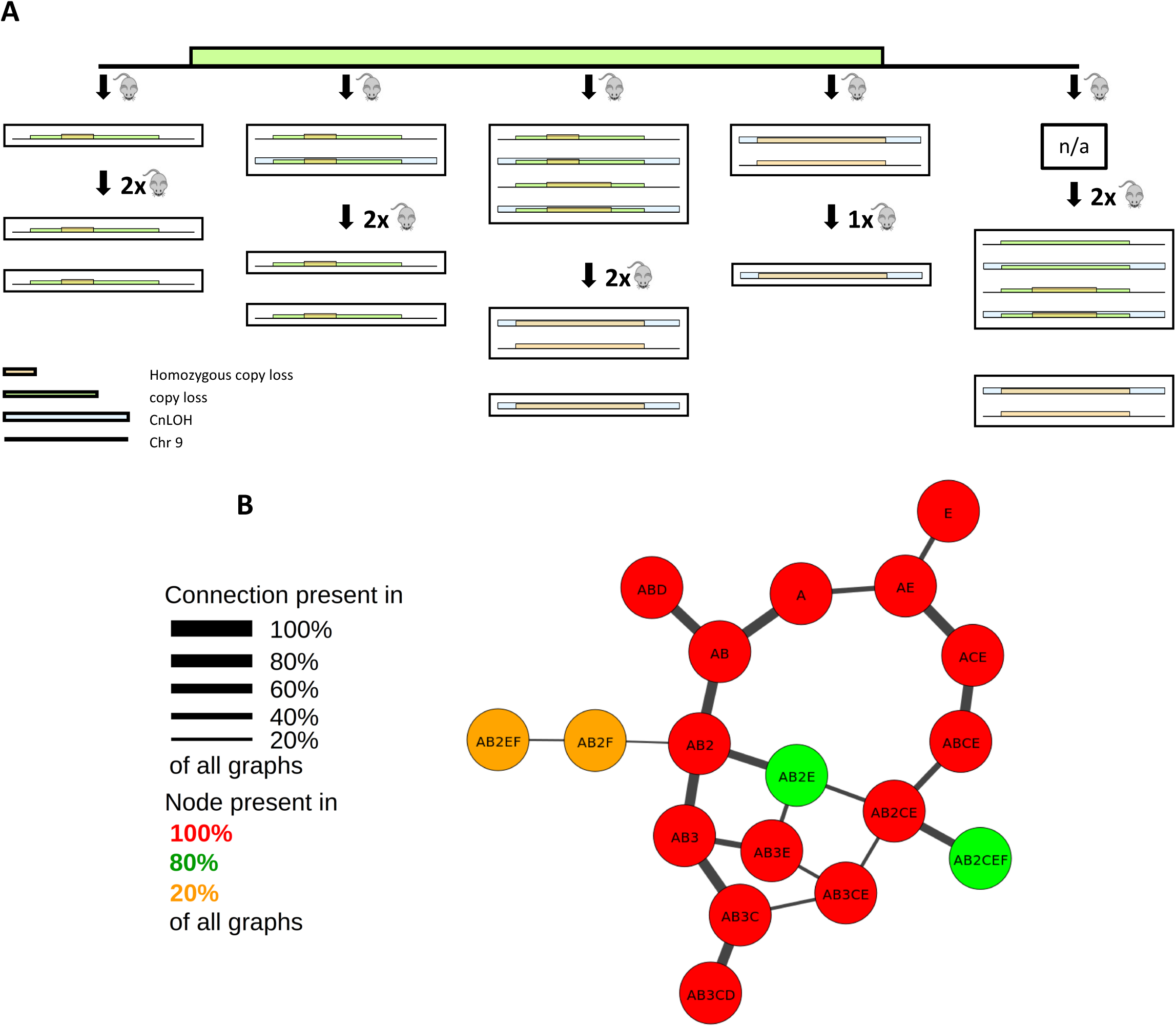
Detailed analysis on chromosome 9 reveals higher clonal complexity (Patient 2). **A.** In depth analysis of CNA on chromosome 9 during serial transplantation suggests a more complex clonal hierarchy with multiple deletions of different size targeting the locus and occurrence of cnLOH. Each box indicates an individual mouse. **B.** Phylogenetic tree of leukemia cells based on SNP-array data obtained at diagnosis and during serial transplantation. Mosaicim of CNA suggested the existence of sub-clones within analyzed samples. The presented tree represents the minimum number of diagnostic clones required to explain the mutational landscape identified in the engrafted leukaemias. Minimal clone numbers were calculated by mathematical modelling (see supplementary materials).

We subsequently sought to shed light on the phenotype-genotype relationship that defines the properties of individual leukemic subclones, by performing SNP analysis on bulk tumour cells and flow-sorted immunophenotypically-distinct ALL subpopulations from three Down’s syndrome ALL patients both before and after xenotransplantation (see schemata in Figure 3A). Analysis of samples from patients 2 and 3 (supplementary Figure 1) revealed identical clonal architectures in diagnostic bulk, stem/B and proB-like cells for both patients (Figure 3BC). All samples analysed from patient 3 confirmed the existence of a single dominant leukemic clone (Figure 3C and supplementary table 1). In addition to the founding chromosome 21 amplification, a third patient (pt.4) we examined harboured a BCR-ABL translocation. FISH analysis showed that in this leukemia the fusion gene marked proB-and preB-like sub-compartments, but not cells with the more primitive and disease-specific stem/B (34^+^38^-^19^+^) phenotype^15^ (Figure 4A). Notably, in serial transplantation experiments flow-sorted stem/B cells only gave rise to grafts with stem/B like cells but not more differentiated (proB-preB) B-cell phenotypes. These stem/B like grafts remained BCR-ABL-negative. Similarly, proB cells derived grafts only contained proB and preB cells, all of which retained the BCR-ABL-fusion. Collectively these data suggest lack of phenotypic plasticity amongst distinct diagnostic cells from this patient.

**Figure 3:**
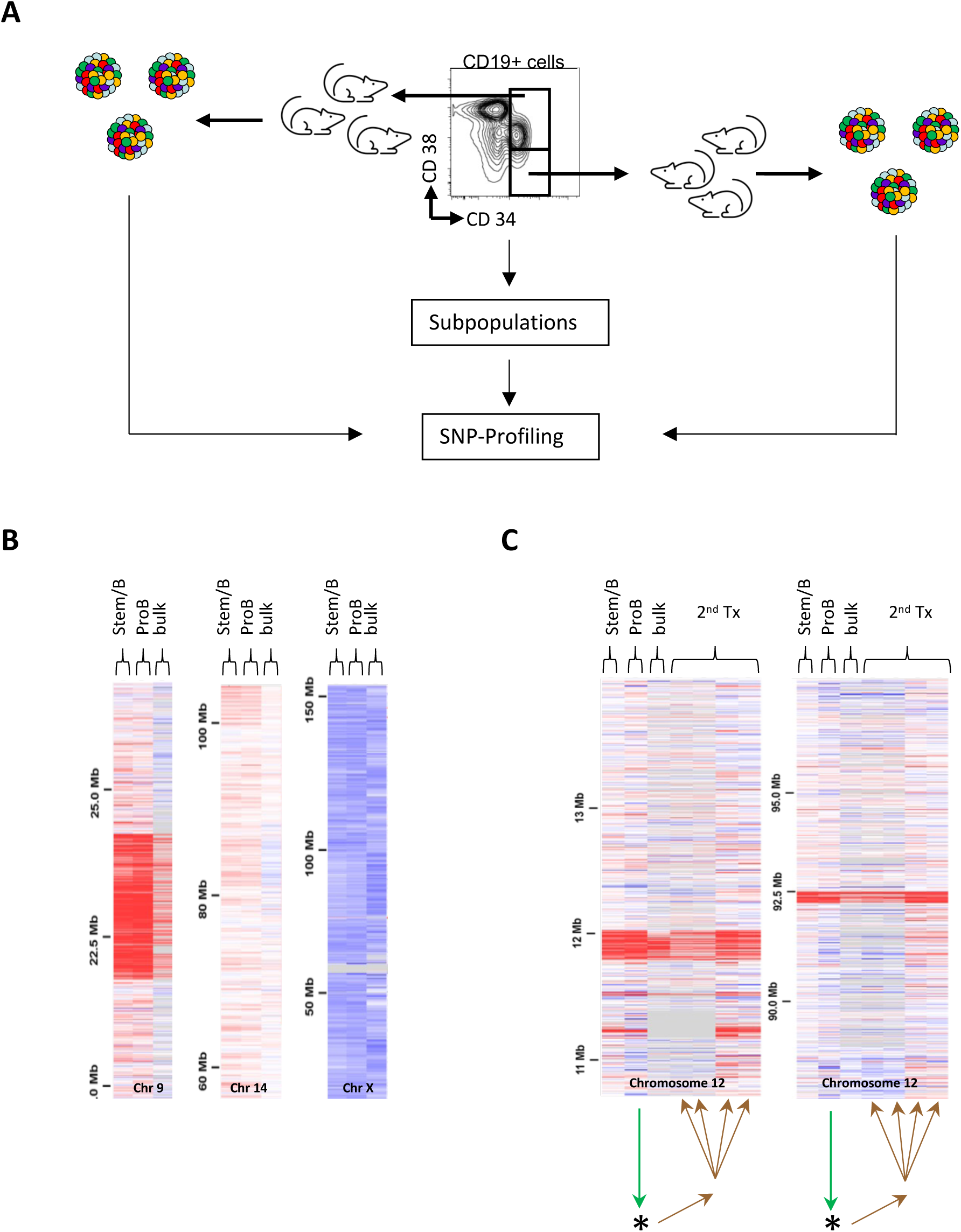
Correlation of phenotype and genotype. **A.** Experimental setup. BM bulk, StemB (CD34^+^CD38^-/low^CD19^+^) and ProB (CD34^+^CD38^+^CD19^+^) cells were analyzed by SNP-arrays before and (if available) after serial transplantation in NSG mice. **B.** In patient 2 bulk, stemB and ProB cells showed identical CNA. **C.** In Patient 3 animals transplanted with bulk, stemB, and ProB cells showed identical CNA profiles: suggesting the existence of a single dominant clone. * Indicate secondary transplants from a primary transplant for which SNP-array data are unavailable due to insufficient numbers of cells for analysis. Secondary transplants from Stem/B and bulk cells did not engraft.

**Figure 4:**
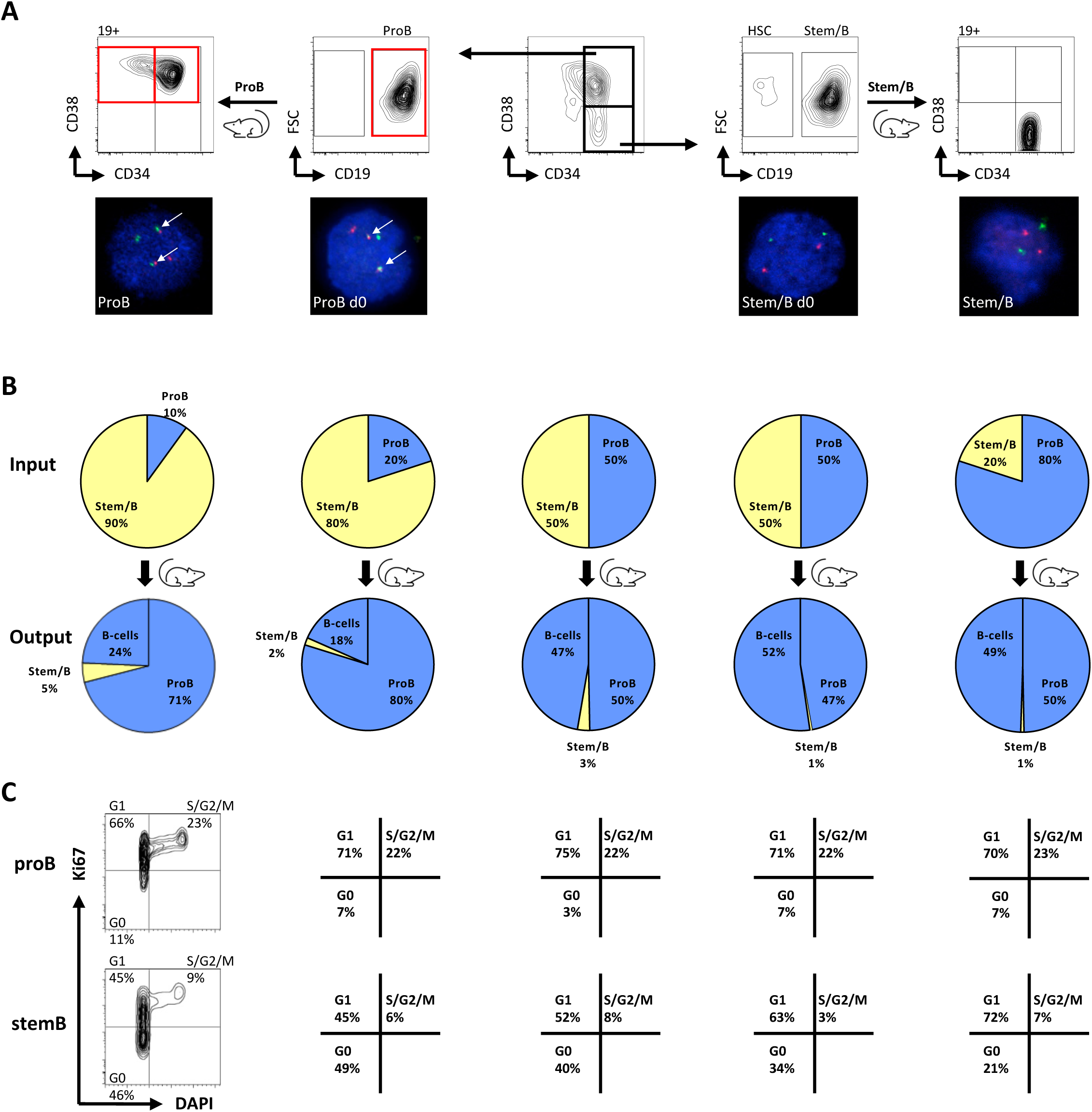
Phenotypically, genetically and functionally distinct sub-clones co-exist in a case of BCR-ABL1^+^ ALL. **A.** FISH analysis of leukemic subpopulations at diagnosis and after serial xenotransplantation in Patient 4 using a BCR/ABL-specific probe. To confirm the leukemic status, sorted cells were analysed by FISH for the appropriate clonal marker. Presence of the t(9;22) translocation is indicated by detection of one green, one red and two red-green fusion signals. **B.** Competitive transplants of genetically and phenotypically distinct BCR-ABL^+^ ProB (CD34^+^CD38^+^CD19^+^) and BCR-ABL^-^ StemB (CD34^+^CD38^-/low^CD19^+^) cells. Input and output ratios of the two populations in individual mice are displayed. **C.** Cell cycle analysis of leukemic ProB (upper panels) and StemB (lower panels) cells from competitively transplanted NSG-mice with subpopulations from a BCR-ABL1^+^ ALL patient. Percentages of cells in G_0_ (DAPI^-^Ki67^-^), G_1_ (DAPI^-^Ki67^+^) and S/G_2/_M (DAPI^+^Ki67^+^) are shown.

We next compared the relative fitness of the two subpopulations (stem/B and proB) in a competitive transplantation setting facilitated by (i) the presence of the BCR-ABL translocation as a clonotypic marker and (ii) the non-interchangeability of their immunophenotypes. We transplanted NSG-mice with different ratios of stem/B and proB-like ALL cells and read out engraftment. In all cases the BCR-ABL^+^ ProB-like cells displayed increased engraftment fitness, most clearly exemplified in the setting where proB cells represented only 10% of the initial inoculum and yet constituted 95% of the resultant leukaemia (Figure 4B). Reasoning that enhanced fitness may stem from a proliferative advantage we next analysed the cell cycle status of engrafted cells. In all engrafted recipients proB cells displayed a more active cycling status compared to their stem/B counterparts, which had a significantly larger proportion of dormant (G_0_) cells (Figure 4C and Supplementary Figure 2).

We subsequently speculated that the different cycling properties of these compartments could be expected to affect their relative sensitivity to therapy. To experimentally determine how these different ALL sub-compartments behaved during therapy, we analysed leukemic subpopulations from serially collected diagnosis, remission, and relapse specimens. ProB-like ALL cells were indeed preferentially eradicated during chemotherapy. Indeed, while both populations were shrank by treatment to similar frequency (0.01%), ProB cells initially represented approximately 70% of leukaemic cells detected at diagnosis while stem/B-like ALL cells were only 8% (Figure 5A). After induction chemotherapy extremely few proB-like cells remained, none of which carried the BCR-ABL translocation. While the behaviour of these subpopulations during treatment contrasts starkly with their fitness as assessed by xenotransplantation, it is entirely consistent with the known relative vulnerability of cycling versus dormant cells to chemotherapy. Similarly, to what has previously been observed in the context of bacteria and antibiotics resistance, these data also support the idea of the existence of a trade-off on phenotypic traits between environments with and without drug^16^. The reduced competitive ability of the less proliferative species (StemB cells) in the absence of treatment hence becomes an advantage in the presence of chemotherapy, while the highly proliferative ProB cells pay a cost to their initial fitness.

**Figure 5:**
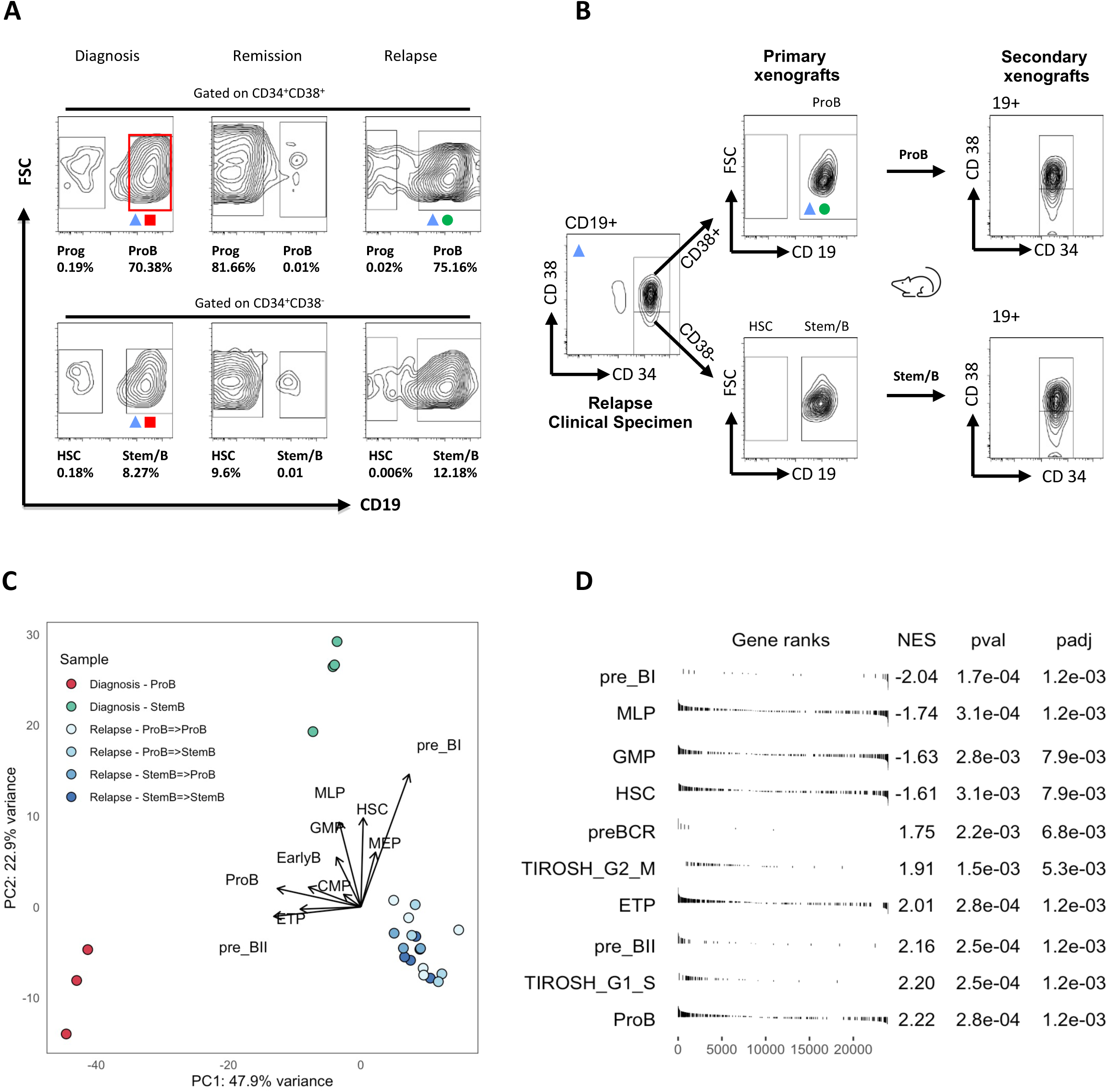
Selective therapy resistance of CD34^+^CD38^-/low^CD19^+^ Stem/B cells in high risk BCR-ABL1^+^ ALL. **A.** FACS analysis of BCR-ABL1 ALL subpopulations at diagnosis, remission, and relapse (patient 4). Percentages of CD34^+^CD38^-/low^CD19^-^ (HSC), CD34^+^CD38^-/low^CD19^+^ (StemB), CD34^+^CD38^+^CD19^+^ (ProB) cells and CD34^+^CD38^+^CD19^-^ (Progenitors) in total bone marrow (BM) mononuclear cells (MNC) are shown. The black boxes show the gates used for frequency calculations. Not all collected events are shown. **B.** FACS analysis of BM cells from serially transplanted NSG mice injected with sorted StemB and ProB cells from patient 4 relapse. **C.** PCA analysis based on bulk RNA sequencing of diagnostic and relapse cells harvested from primary and secondary xenografts. Six different populations of cells were analysed. i) diagnostic stemB cells, ii) diagnostic proB cells iii) relapse stemB cells harvested from secondary xenografts injected with stemB cells isolated from primary xenografts iv) relapse proB cells harvested from secondary xenografts injected with stemB cells isolated from primary xenografts, v) relapse proB cells harvested from secondary xenografts injected with proB cells isolated from primary xenografts, and vi) relapse stemB cells harvested from secondary xenografts injected with proB cells isolated from primary xenografts. Arrows showing the contribution of different gene expression signatures (calculated as per panel D) to the sample’s clusters biology are shown. **D.** Gene set enrichment analysis (GSEA) for a selection of MSigDB-50 Hallmark gene-sets and published signatures associated with predefined differentiation stages of the B-cell ontogeny^19^. ProB diagnostic cells are compared to the stemB diagnostic cells. Significance thresholds: <0.05 after adjusting for multiple testing (using the Benjamini-Hochberg method).

Interestingly, the relapse sample was populated by both stemB and proB-like cells (Figure 5A). However, in contrast to the situation at diagnosis leukemic proB cells present at relapse were BCR-ABL-negative and both relapse stemB and proB cells showed phenotypic plasticity upon serial transplantation (Figure 5B). This data is somewhat counterintuitive in that it argues against the simplest origins of relapse from the predominant stemB or proB clones present at diagnosis. The stem/B compartment at presentation lacked phenotypic plasticity in that it was unable to regenerate proB- or preB-like cells. Since the proB cells at relapse were BCR-ABL-negative, relapse could also not have been seeded by the predominant diagnostic proB-like compartment that was positive for BCR-ABL.

Taking advantage of the xenograft system we further investigated the phenotypic properties of diagnostic and relapse cells at the molecular level by performing small cell number bulk RNA sequencing (RNAseq) on cells harvested from either primary and or secondary xenografts. We hence characterised six distinct cell populations i) diagnostic stemB cells from primary xenografts, ii) diagnostic proB cells from primary xenografts iii) relapse stemB cells harvested from secondary xenografts injected with stemB cells isolated from primary xenografts iv) relapse proB cells harvested from secondary xenografts injected with stemB cells isolated from primary xenografts, v) relapse proB cells harvested from secondary xenografts injected with proB cells isolated from primary xenografts, and vi) relapse stemB cells harvested from secondary xenografts injected with proB cells isolated from primary xenografts. Principal Component Analysis (PCA) showed clustering of samples based on disease stage. Interestingly, diagnostic cells also clustered separately based on differentiation state, while all relapse populations clustered closely together (Figure 5C). This finding resonates well with our observation that in this patient leukaemia, relapse but not diagnostic cells displayed high cell plasticity. While the PC1 component of the PC captured most of the sample variance (47.15), PC2 clearly separated diagnostic proB cells and all relapse subpopulations from diagnostic stemB cells. Pathways analysis on the genes driving this PC component revealed enrichment for TNF-alpha via NF-KB signalling cascade, which has previously been implicated in regulation of cell survival, proliferation and stemness^17^.

Gene set enrichment analysis (GSEA) using publicly available gene expression signatures^18,19^ also validated our FACS analysis results in that compared to diagnostic proB cells and relapse cells, diagnostic stemB cells downregulated cell proliferation associated genes and expressed higher levels of markers typical of more immature stages of the haematopoietic and B-cell differentiation hierarchy (such as HSC and MLP cells) (Figure 5C-D and Supplementary Figure 3A-B). Most B-lineage cells from ALL patients are thought to be arrested at some point of the transition from the proB to preB stage, therefore broadly allowing the classification of ALL leukaemias into two distinct subtypes based on preBCR function. Those arrested at the preBI stage rely upon IL7R/STAT5 signaling (preBCR^-^), while those arrested at preBII stage signal by the preBCR (preBCR^+^)^20^. The frequency of pre-BCR^+^ or pre-BCR^-^ cases has also been suggested to associate with different cytogenetic subtypes of childhood ALL. For example, the majority of BCR-ABL1^+^ leukaemias tend to display constitutive active cytokine signalling via activation of STAT5 and repression of BCL6. Herein, however, we found that these two genetically and phenotypically distinct disease subsets can coexist within the same leukaemia. Interestingly, contrary to expectations, diagnostic proB cells, which carried the BCR-ABL translocation and expressed markers associated with later stages of the B-cell differentiation (Figure 5C), displayed stronger expression of the preBCR receptor components and signalling cascade, while the BCR-ABL^-^ diagnostic stemB cells preferentially upregulated various genes involved in the STAT5 signalling pathway (Supplementary Figure 3D).

We subsequently used SNP arrays to genetically characterise different phenotypic compartments at presentation, relapse and after serial transplantation and track the phylogenetic origins of the relapse propagating clones (Figure 6). SNP analysis of the relapse samples indicated that it lacked several of the distinctive genetic abnormalities which characterised either the dominant stemB compartment at diagnosis or the dominant proB. However, the relapse hierarchy shared with the diagnostic compartments deletions on both chromosomes 12 and 14. Of note, diagnosis and relapse clones also shared a common VDJ rearrangement (data not shown).

**Figure 6:**
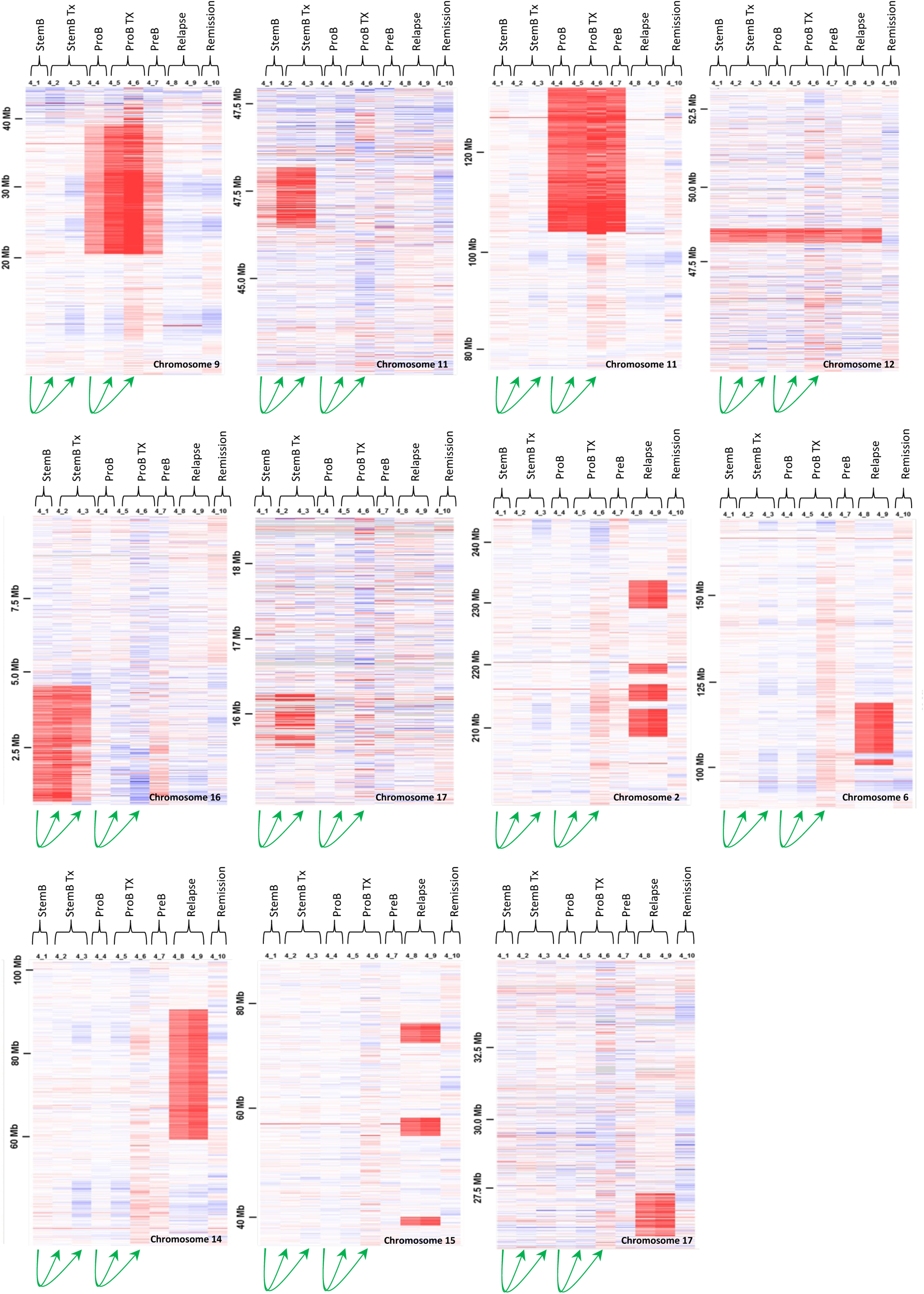
Distinct sub-clones in the BCR-ABL1^+^ ALL and the BCR-ABL^-^ relapse clones are genetically related (Patient 4). **A.** SNP array results of diagnostic stem/B, xenografted stem/B, diagnostic proB, xenografted proB, diagnostic preB, relapse StemB, relapse ProB and MRD-negative Remission sample are displayed. A chromosome 12 deletion was present in all but the remission control sample illustrating the clonal relationship of all leukemic samples. CNA of diagnostic stem/B and ProB were identified and stable during serial transplantation with identification of some mosaicism suggesting sub-clones in each compartment. Relationships of transplants are indicated. Relapse stem/B and proB were genetically identical but stem/B showed 90% mosaicism of all clones suggesting the existence of a pre-leukemic clone. **B.** Detailed analyses SNP array revealed different sizes of some chromosome deletions at diagnosis, remission and relapse and suggest the existence of distinct pre-leukemic sub-clones for which a clonal hierarchy cand be re-constructed.

**Figure 7:**
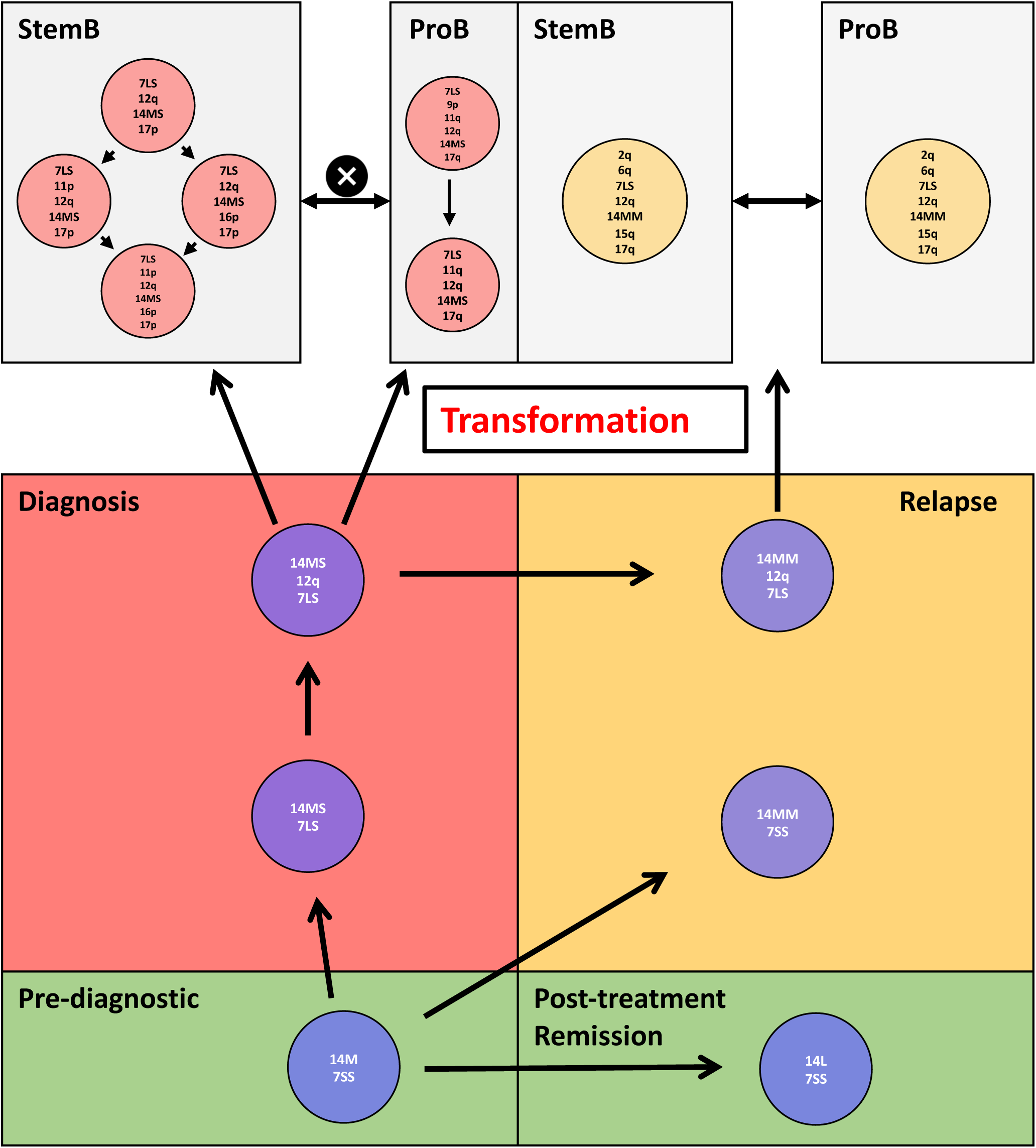
Pre-leukemic and leukemic hierarchy. Based on genetic mosaicism and different sizes of chromosome 7 and 14 deletions (Patient 4) a pre-leukemic clonal hierarchy could be constructed which showed clonal evolution at the pre-leukemic stage that gave rise to the diagnostic leukemic sub-clones as well as evolving into another pre-leukemic sub-clone responsible for relapse.

Altogether, this data implies the survival over treatment of a common evolutionary ancestor not detectable at diagnosis which gave rise to the relapse hierarchy through further diversification. Accordingly, analysis of a bulk remission sample identified mosaic deletion of chromosome 14 in the absence of del12 and any other obvious genetic abnormality. Therefore, suggesting that this mosaic deletion is not a germline event and, likely, the earliest genomic alteration (also preceding del12). To understand the basis of the mosaicism observed in the remission sample, we separated CD3+ and CD3-cells and performed SNP analysis. The CD3-compartment contained no evidence of the 14del. In contrast the CD3+ cells showed a complex pattern of chromosome 14 deletions indicative of a progressive multi-step evolutionary process (Supplementary table 2). We next traced these 14del variants back to the diagnostic and relapse stemB and proB compartments, as well as the CD3+ compartment isolated at diagnosis. This spectrum of 14 deletions provides a branched framework for subsequent evolution of stemB and proB clones seen at diagnosis and relapse. Thus, using chromosomal deletions as markers of lineal ancestry we have here reconstructed the evolutionary history of a Down’s ALL from the predisposing event (Trisomy 21), through the acquisition of initiating lesions giving rise to a preleukemic hierarchy, into further transformation marked by the onset of frank leukaemia. We also monitored the response of this complex leukemic hierarchy to chemotherapy through the analysis of matching remission and relapse specimens. Beyond the genetic considerations our work emphasizes the role of cellular state primarily at the level of dormancy and cell cycle indicating vulnerability to chemotherapy.

## Discussion

Many studies have explored the extent and impact of different, individually assessed, sources of intratumor heterogeneity on the evolution of tumours (particularly under treatment). Our work, however, argues that cancer should be viewed as a complex matrix of genetic and epigenetic heterogeneity such as cellular phenotype and cell cycle properties which, altogether rather than in isolation, provide a substrate for tumour evolution during disease progression in response to the selective impetus of therapeutic intervention.

The idea that epigenetically determined cell states, contribute to functional heterogeneity amongst tumour cells by controlling fundamental cellular properties such as, for example, cell identity, phenotype and cell-cycle stage is paradigmatically exemplified by the cancer stem cell (CSC) hypothesis; which postulates that cells within a tumour are organized in a hierarchical fashion reflecting lineage relationships and tumorigenic potential, and that the maintenance of cancer clones is uniquely dependent on the most primitive CSCs with self-renewal capacity^21–23^. This notion, which was pioneered in the context of blood malignancies, remains however extensively debated. While phenotypic heterogeneity - primarily at the level of immune-phenotype – has been widely observed in ALL, disparate data exist concerning the existence of LSCs within this disease. Earlier work by Rehe et al., showed that many cellular fractions contain LSC-potential at a similar frequency and that the majority of - if not all - cells may be able to propagate the disease^24^. Importantly, the same study also found evidence for phenotypic plasticity within different subpopulations upon serial transplantation in immune-deficient mice. Distinct leukaemia stem cells (LSCs) have nonetheless been identified in acute myeloid leukaemia (AML), where functional evidence for their existence has been provided^25–28^. However, even in this context, their exact phenotype remains elusive, with more recent studies uncovering LSC activity across different cellular subsets^29^. We here provide evidence that although different Down’s ALL disease compartments can all contribute to leukaemia propagation *in vivo,* these same compartments display largely distinct fitness; both in the presence and in the absence of chemotherapy. In this scenario, more primitive stem-like compartments are outcompeted by highly proliferative early-B cell progenitors in the absence of treatment. However, this selective advantage is reversed in the presence of chemotherapy, which selects for quiescent cells. While the genetically heterogenous nature of childhood ALL has been demonstrated by multiple earlier studies, our current work provides the first direct evidence for genetic diversity of cancer propagating cells in patients with Down’s syndrome ALL. Our SNP array analysis of serially transplanted ALLs shows that leukemic cells undergo a process of gradual branching clonal evolution demonstrated by the coexistence within a leukaemia of multiple subclones bearing alterations of the same genomic locus characterised by distinct breakpoints.

Our earlier work showed that in many ALL leukemias genotype and phenotypes with relevance to chemoresistance do not segregate. As a result, induction chemotherapy does not select at the level of genotype. We here identified and studied the evolutionary history of an unusual case of BCR-ABL+ Down’s ALL in which genotype and phenotype do instead clearly segregate. In this patient’s leukaemia the stem/B cells were fusion-gene-negative while proB/pre-B cells were fusion-gene-positive. This case allowed us to reconstruct in detail the complex genotype-phenotype relationship characterizing different leukemic cell fractions labelled by both distinct genetic makeup and fitness. We hence observed that highly quiescent and developmentally more primitive cells preferentially escaped treatment. As these cells carried only a subset of diagnostic genetic alterations, the relapse disease was genetically divergent from the diagnostic specimen. Reconstruction of the leukaemia phylogenetic tree demonstrated that the dominant population at relapse originated from a rare pre-leukemic clone. This was likely present at diagnosis below detection level and, in line with its likely low proliferative activity, did not extensively expand upon transplantation.

We first demonstrated the existence of pre-leukemic cells in ALL in studies of mono-chorionic twins of whom one was diagnosed with TEL-AML1-positive ALL^15, 30^. Blood spot studies from Guthrie-cards further showed that these pre-leukemic cells can precede overt leukaemia by up to 12 years and are also detectable in patients that never go on to overt disease^31, 32^. A case report of a 7-year old boy diagnosed with ALL with a mixed lympho-myeloid phenotype previously suggested that BCR-ABL+ non-Down leukaemic cells could reactivate to drive disease recurrence more than two decades from initial diagnosis^33^. Our new data add to our understanding of this elusive population by showing that Down Syndrome ALL chr21+ cells, which have not yet acquired the BCR-ABL translocation labelling the more proliferative proB compartment disease, can preferentially survive chemotherapy and, therefore, represent a reservoir for relapse initiation. Whether these cells are quiescent principally as a function of their unique developmental origin^34^ or rather of their genotype^35^ or a combination of both remains to be investigated and would be important to understand if new therapeutic interventions targeting these cells are to be designed. This notwithstanding, pre-leukemic clones, even when rare, should clearly be considered part of the fully transformed leukaemia landscape and may upon selection by therapy provide an additional cellular source for subsequent relapse.

## Acknowledgments

We are grateful for the support of Leukaemia and Lymphoma Research (TE; Specialist Programme Grant), Cancerfonden (TE; Project grants CAN 2013/712 and 20 1334 PjF), MRC Molecular Haematology Unit core award (SEJ; MC_UU_12009/5), Blood Cancer UK (SEJ), Swedish Childhood Cancer Fund (SEJ), the National Institute for Health Research Biomedical Research Centre Programme, the CBRC, UCL, Haematolinne, the Leukemia and Lymphoma Society, USA (Career Development Program fellowship to PSW), the Deutsche Forschungsgemeinschaft (fellowship to CL; LU1474/1-1 and programme grant as part of the Sonderforschungsbereich; SFB873, Teilprojekt A13).

## Authorship Contributions

CL and VT performed experiments; RC analysed the SNP array; JH analysed the RNAseq data with input from VT; AS interpreted and supervised SNP-array data analyses; TS developed the mathematical model; CL and PSW performed the cell cycle analysis, VB carried out FISH analyses; AC provided patient data and samples; SAC, HF and PSW performed cell sorting. CL, VT, SEWJ and TE designed the study and interpreted data. CL, VT and TE wrote the paper with critical input from other authors.

---

**Figure S1:**
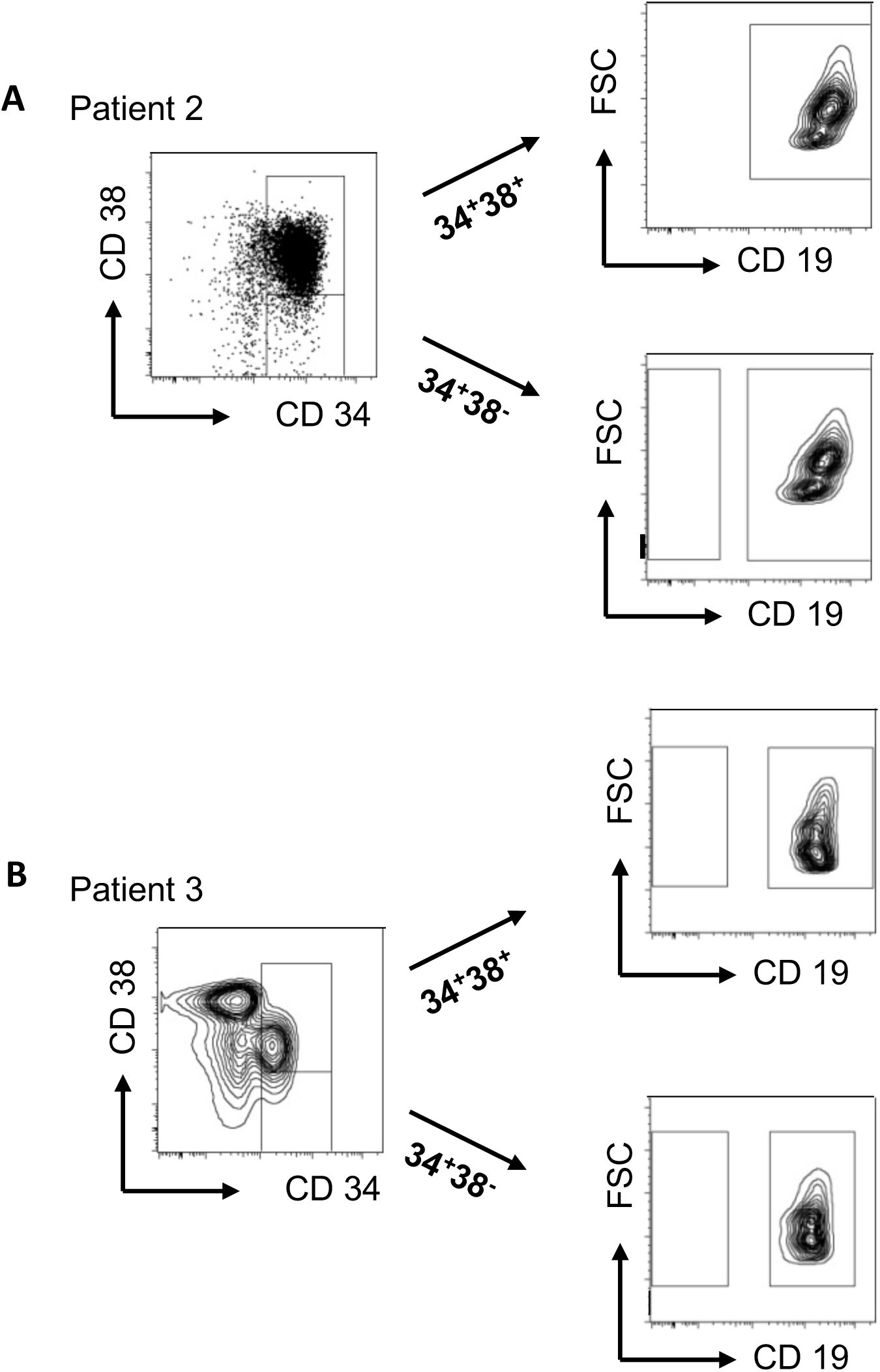
Flow cytometry sorting strategy. Representative gating strategy used for the analysis of immunophenotypically-distinct Down’s syndrome ALL subpopulations before and after xenotransplantation. StemB (CD34^+^CD38^-/low^CD19^+^) and proB (CD34^+^CD38^+^CD19^+^) cells were sorted. **A-B**. Patient 2 and 3 leukaemia respectively.

**Figure S2:**
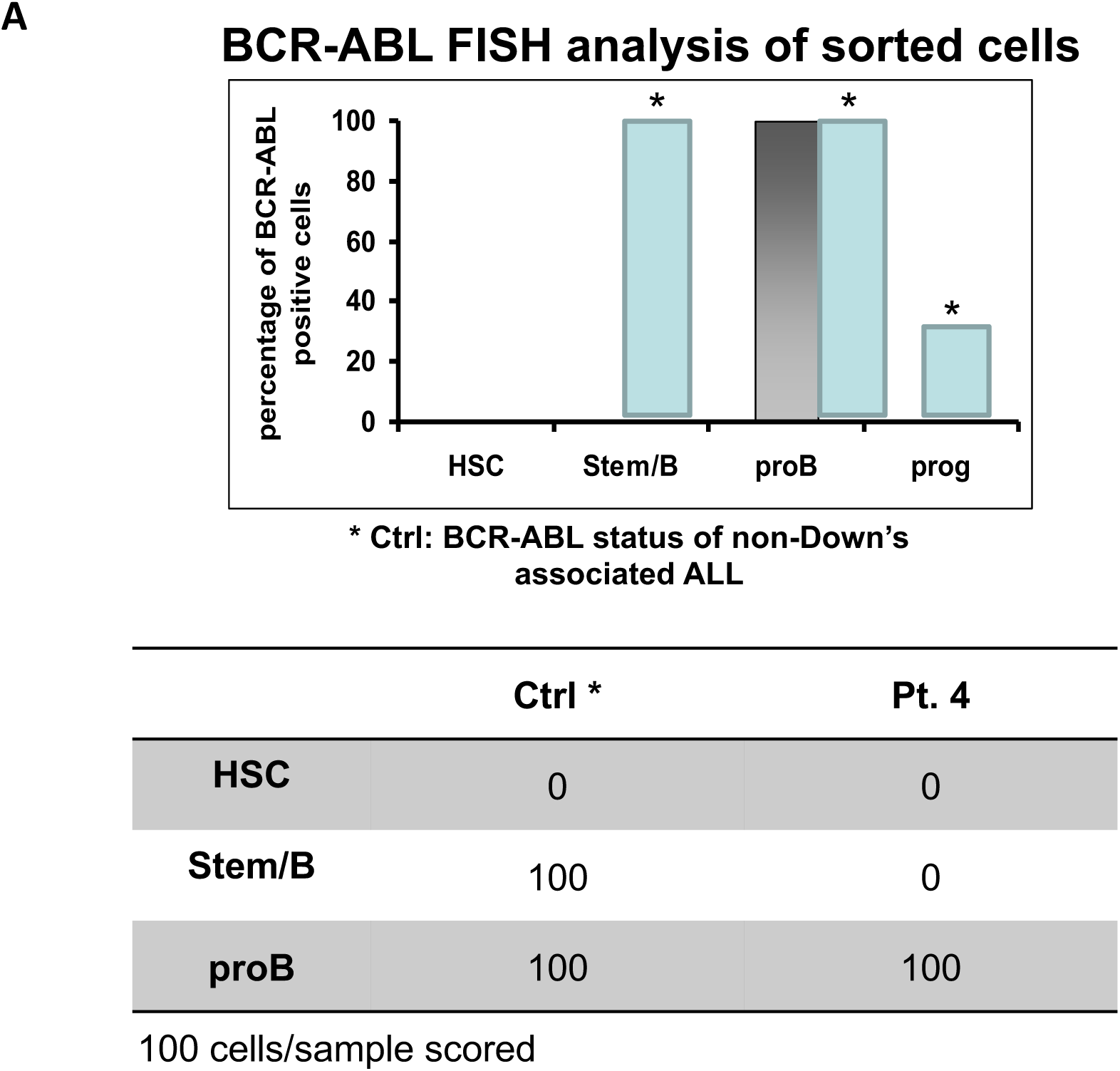
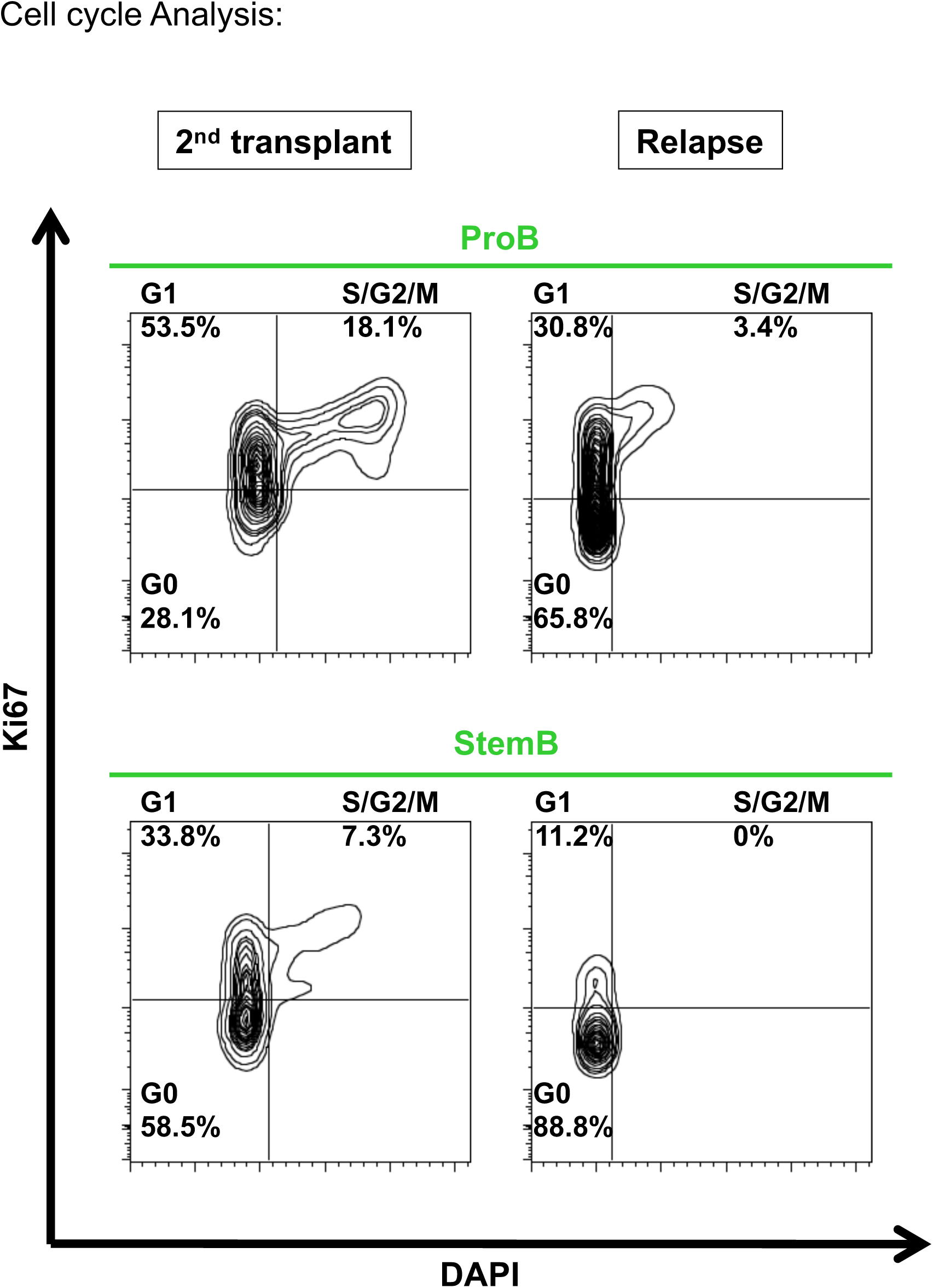
Cell cycle analysis. **A.** Cell cycle analysis of representative xenografted stemB and ProB as compared to the **B.** patient relapse.

**Figure S3:**
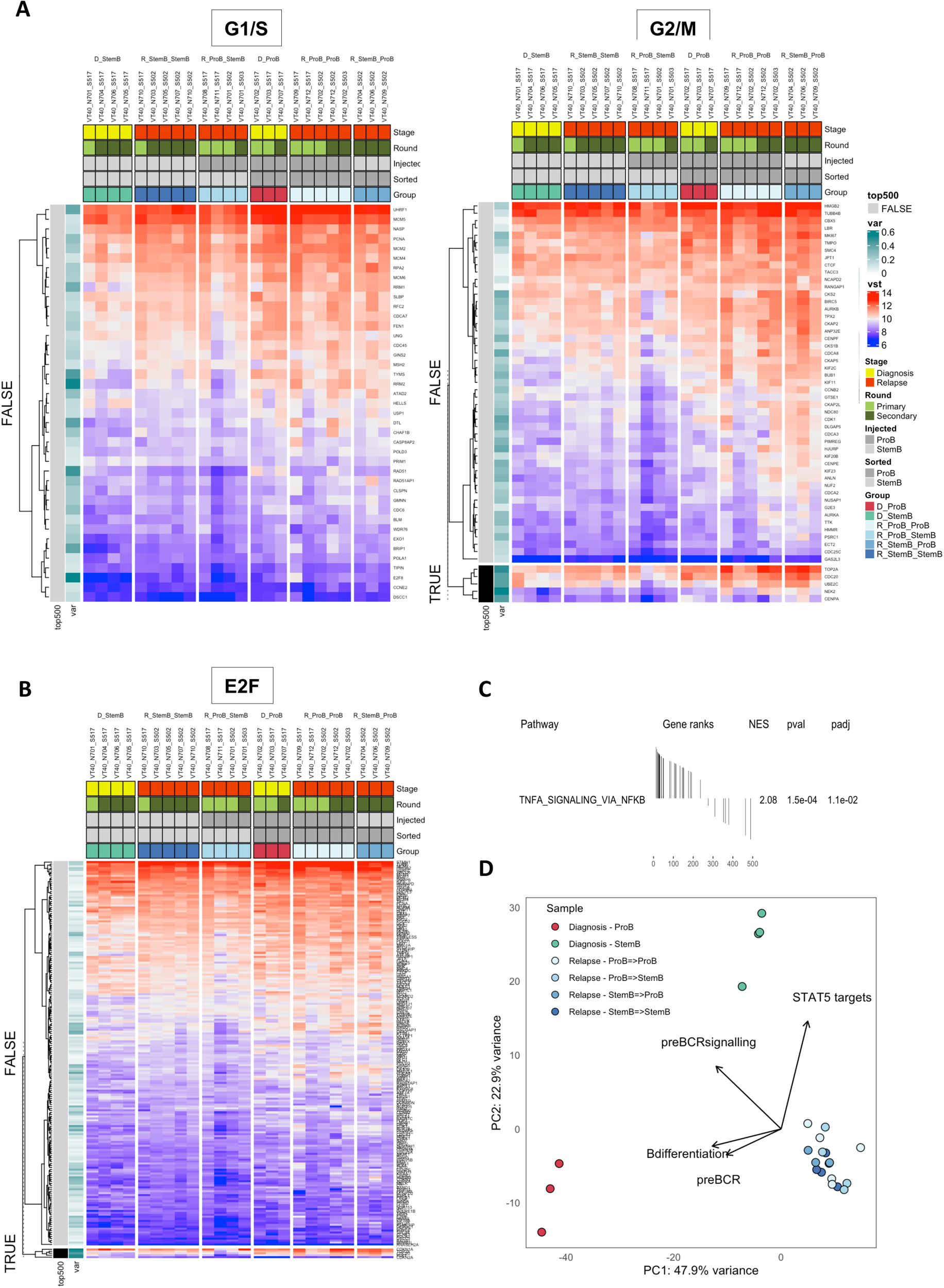
RNA sequencing analysis. **A-C** Gene Set Variation Analysis (GSVA) analysis of RNA sequencing data looking at the expression of cell cycle related gene signatures. Six different populations of cells were analysed: i) diagnostic stemB cells, ii) diagnostic proB cells iii) relapse stemB cells harvested from secondary xenografts injected with stemB cells isolated from primary xenografts iv) relapse proB cells harvested from secondary xenografts injected with stemB cells isolated from primary xenografts, v) relapse proB cells harvested from secondary xenografts injected with proB cells isolated from primary xenografts, and vi) relapse stemB cells harvested from secondary xenografts injected with proB cells isolated from primary xenografts. Within each heatmap samples are sorted by category and genes split into top 500 most variable vs all others. **A.** G1/S gene signature^36^, **B.** G2/M gene signature^36^, **C.** E2F targets gene signature. **C.** Gene set enrichment analysis (GSEA) for a selection of MSigDB-50 Hallmark gene-sets and published signatures associated with predefined differentiation stages of the B-cell ontogeny^19^. This analysis looks specifically at the top 500 genes driving clustering along the PC2 component of the PCA analysis. **D.** PCA analysis based on bulk RNA sequencing of diagnostic and relapse cells harvested from primary and secondary xenografts. Arrows showing the contribution of gene expression signatures involved in BCR signalling and B-cell differentiation are shown. Significance thresholds: <0.05 after adjusting for multiple testing (using the Benjamini-Hochberg method).

